# Dnmt1a is essential for gene body methylation and the regulation of zygotic genome activation in the wasp

**DOI:** 10.1101/2021.02.02.429402

**Authors:** Deanna Arsala, Xin Wu, Soojin V. Yi, Jeremy A. Lynch

**Author notes:** Corresponding author; Jeremy A. Lynch.

## Abstract

Gene body methylation (GBM) is an ancestral aspect of DNA methylation (Sarda, Zeng, Hunt, & Yi, 2012; Yi, 2012; Zemach, McDaniel, Silva, & Zilberman, 2010) whose role in development has been obscured by the more prominent roles of promoter and CpG island methylation. The wasp Nasonia has little promoter and CpG island methylation, yet retains strong GBM (Park et al., 2011; Wang et al., 2013; Werren et al., 2010), making it an excellent model for elucidating the role of GBM. Here we show that Nasonia DNA methyl transferase 1a (Nv-Dnmt1a) knockdown leads to failures in cellularization and gastrulation of the embryo. Both of these disrupted events are hallmarks of the maternal-zygotic transition (MZT) in insects. Analysis of the embryonic transcriptome and methylome revealed strong reduction of GBM and widespread disruption of gene expression during embryogenesis after Nv-Dnmt1a knockdown. There was a strong correlation between loss of GBM and reduced gene expression in thousands of methylated loci, while affected unmethylated genes tended to be upregulated. We propose that reduced GBM and subsequent lower expression levels of methylated genes was the direct effect of Nv-Dnmt1 knockdown, and that this disruption led to widespread downstream dysregulation of MZT, and manifesting in developmental failure at gastrulation.

**Significance Statement:** The importance of gene-body methylation (GBM) in development is unclear, due to the difficulty in teasing apart the effects of cis-regulatory methylation from those of GBM in vertebrate model systems. Unlike vertebrate models, the methylation machinery in the jewel wasp *Nasonia vitripennis* appears to exclusively mediate GBM, thus simplifying interpretation of the role of GBM in development. Knockdown of DNMT1 (Nv-Dnmt1a) in *Nasonia* leads to embryonic lethality, which we show is caused by a failure of cellularization and gastrulation. Nv-Dnmt1a knockdown resulted in a global loss of GBM in the embryo, which was strongly correlated with a down-regulation of gene expression. We propose that GBM facilitated by Nv-Dnmt1a is required for proper zygotic genome activation in the wasp.

## Introduction

The maternal-zygotic transition (MZT) is an essential stage in multicellular eukaryotic development and represents the transition of developmental control from maternal genome products provided to the egg during oogenesis to the transcriptional products of newly formed zygotic genome (1, 2).

The MZT is regulated by many layers of pre- and post-transcriptional mechanisms, including histone modification, chromatin packaging, miRNAs, mRNA decay and degradation, and methylation of genomic DNA (1,2). The interactions between the maternal and zygotic genomes have been most comprehensively demonstrated in the insect model *Drosophila melanogaster*. Maternal transcript destabilization, which is required for zygotic genome activation (ZGA), is carried out in part by maternally deposited mRNAs encoding RNA binding proteins (RBPs) (3, 4) and aided by transcription factors such as Zelda (Zld), which promotes miRNA expression that further destabilize maternal transcripts. Under normal circumstances Zld also increases chromatin accessibility and helps activate thousands of zygotic genes to their proper level (5–7). Disruption of either zygotic genome activation, or maternal transcript clearance leads to embryonic lethality. In both cases, death is due to failures in the earliest developmental events that are dependent on the zygotic genome: cellularization of the syncytial blastoderm, and gastrulation (5, 8).

Most of the strategies for regulating the *D. melanogaster* ZGA (e.g., histone modification, chromatin packaging, action of miRNAs, mRNA degradation), and many of the molecules involved, are conserved throughout the animals (9). However, *D. melanogaster* lacks a major class of genomic regulation, in the form of methylation of cytosines in CpG dinucleotides (CpG methylation) (10, 11). CpG methylation can be catalyzed by DNA methyltransferases 1 and 3 orthologs (DNMT1 and DNMT3), is crucial for regulating gene expression in most complex eukaryotes (12, 13) (14), and has a significant role in regulating the MZT in many species (15, 16).

The most well understood functional impact of CpG methylation is in silencing gene expression via methylation of cis-regulatory sequences in clusters of CpG nucleotides (CpG islands) (13, 17). However, it is becoming clear that this is not a ubiquitous regulatory mechanism (18). A more universal, and likely ancestral (19, 20), form of DNA methylation occurs between the transcription start site and transcription end site of a gene, and is known as gene body methylation (GBM) (21).

Previous studies have correlated GBM with less variable, and higher levels of gene expression than genes without GBM (20, 22–25), but its role in regulating gene expression in specific developmental contexts such as the MZT is poorly understood. Whole-genome bisulfite sequencing (WGBS) studies have shown that DNA methylation patterns are dynamic during the MZT in vertebrate model systems (26–28). However, since manipulating DNA methylation machinery in vertebrate models affects both CpG islands and GBM, it is difficult to identify functions of GBM that are independent of CpG island methylation.

In contrast to vertebrates, DNA methylation occurs almost exclusively at gene bodies in insects (29–32). Thus, such insect model systems have the potential to make significant contributions in understanding the role of GBM in development. A particularly attractive system with a full methylation toolkit (harboring orthologs of both DNMT1 and DNMT3) and exclusively gene body methylation is the wasp *Nasonia vitripennis* (30, 33, 34).

There are three DNMT1 paralogs and one DNMT3 ortholog in the *N. vitripennis* genome, and one of them, *Nv-DNA methyltransferase 1a* (*Nv-Dnmt1a*), was shown to be essential for embryogenesis, while knockdown of the other two DNMT1 paralogs and DNMT3 ortholog produced viable embryos (35). Therefore, *N. vitripennis* offers an exciting opportunity to functionally study the specific role of gene body methylation in a well-defined developmental system. In this work, we set out to experimentally investigate the role of *Nv-Dnmt1a* and infer the role for gene body methylation in early embryogenesis of *N. vitripennis*.

Our detailed analysis of effects of *Nv-Dnmt1a* depletion on early embryogenesis revealed major defects in cellularization and morphogenesis of the early embryo, which are phenotypes typically seen after disruption of the MZT in insects. Using parental RNAi (pRNAi) and WGBS we found that the vast majority of gene body methylation is lost when *Nv-Dnmt1a* is knocked down. We then show that loss of gene body methylation during zygotic genome activation is significantly correlated with reduced gene expression from the affected loci during zygotic genome activation. We propose that disruption of methylated gene expression after *Nv-Dnmt1a* knockdown has cascading effects that dysregulate thousands of genes during the MZT, leading to failure in the first developmental processes that rely on a properly activated zygotic genome.

## Results

### Nv-Dnmt1a function is required for early embryonic events that require proper regulation of the zygotic genome

The initial characterization of *Nv-Dnmt1a* showed that knockdown through parental RNAi (pRNAi), where female embryos are injected with dsRNA (36), led to lethality of embryos produced by injected females (35). To gain more insight into the specific functional importance of *Nv-Dnmt1a* for early embryonic development, we examined the embryonic developmental failures following *Nv-Dnmt1a* pRNAi in more detail.

Early development is very rapid in *N. vitripennis*. Nuclei divide rapidly, simultaneously, and without the formation of cell membranes to fill the large, unicellular egg with syncytial nuclei. The first 7 division cycles occur deep in the yolk, after which nuclei migrate to the cortex of the embryo, forming a syncytial blastoderm (37) consisting of a single layer of nuclei populating the entire cortex of the egg. Syncytial blastoderm nuclei divide five more times, prior to cellularization. We observed no differences between control embryos and *Nv-Dnmt1a* pRNAi embryos during the first 11 syncytial divisions (Fig. 1A).

**Figure 1.**
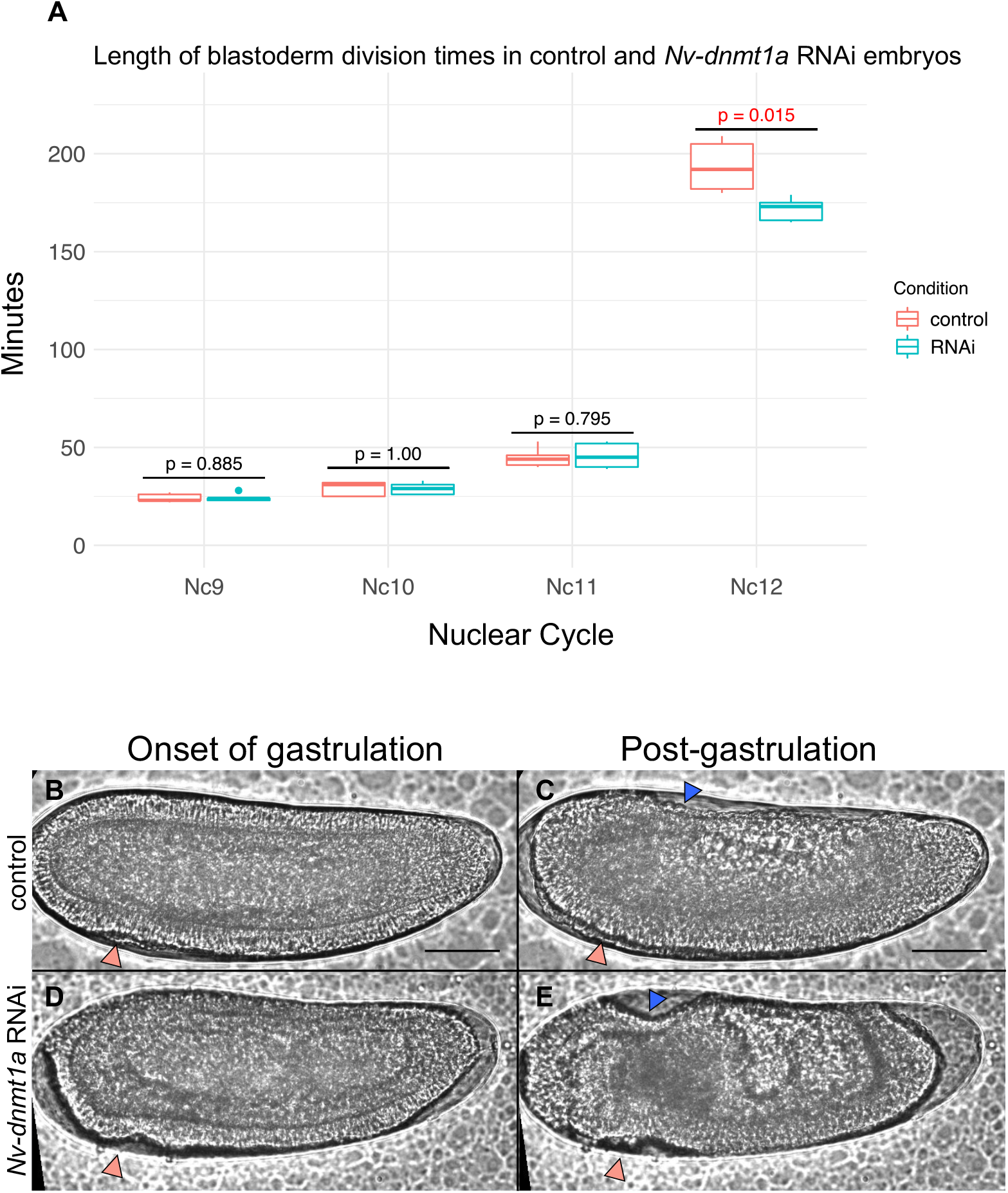
Live imaging of control and *Nv-Dnmt1a* RNAi embryos. A-A’) Live-imaging of a control embryo at the onset of gastrulation and at post-gastrulation. Peach arrows in (A) point to the initial mesodermal folds which are eventually internalized by A’. The blue arrowhead in A’ points to the serosa. Scale bars (black lines in A-A’) are 50μm. (B-B’) Live-imaging of a *Nv-Dnmt1a* RNAi embryo at the onset of gastrulation and at what is equivalent to post-gastrulation. Defective mesodermal folds are denoted by peach arrowheads in B and B’. Abnormal serosal movements are denoted by a blue arrowhead in B’. C) Box and whisker plot displaying the length of blastoderm division times that were measured during live imaging experiments. Nc = nuclear cycle #. P values were determined by performing a 2-tailed T-test assuming unequal variance between RNAi and control. N=5 control, N=5 *Nv-Dnmt1a* RNAi. All scale bars (black lines in A-A’) represent 50 μm.

The first obvious defect occurred at the 12^th^ division cycle (Fig. 1B, 1D). Normally, nuclei become cellularized by encroaching plasma membrane, and immediately begin the morphogenetic movements of gastrulation at the end of this stage (37). The first evidence of morphogenetic movements in control embryos was observed 193 minutes after the onset of nuclear cycle 12, on average (n=5, SD = 13.12, (Fig. 1A)), similar to our previously published results in untreated embryos (38). However, apparent morphogenetic movements in *Nv-Dnmt1a* pRNAi embryos began significantly earlier (171 minutes (n=5, SD = 5.98) into cycle 12, on average) (Fig. 1A). In addition to being premature, these movements were abnormal. In *Nv-Dnmt1a* pRNAi embryos, a line of cells at the anterior end of the embryo began to ingress posteriorly (Fig. 1D) which was not seen in control embryos (Fig. 1B). In addition, internalization of the mesoderm and migration of the serosa from the dorsal pole to cover the embryo both failed (Fig. 1C, 1E, see Buchta et al., for more description of these movements (37)).

To get a more detailed understanding of the cellular basis of these defects, we examined high-resolution images of transverse sections *Nv-Dnmt1a* pRNAi and control embryos at the end of the 12^th^ cell cycle, when the cellularization and the first gastrulation movements should be occurring.

All control embryos in the process of cellularization, displayed membrane ingression that was consistent around the entire blastoderm circumference (Fig. 2A). In contrast, membrane ingression was uneven in *Nv-Dnmt1a* pRNAi embryos, leaving many nuclei completely or mostly unencapsulated prior to gastrulation (Fig. 2C, white arrows point to examples of non-cellularized nuclei).

**Figure 2.**
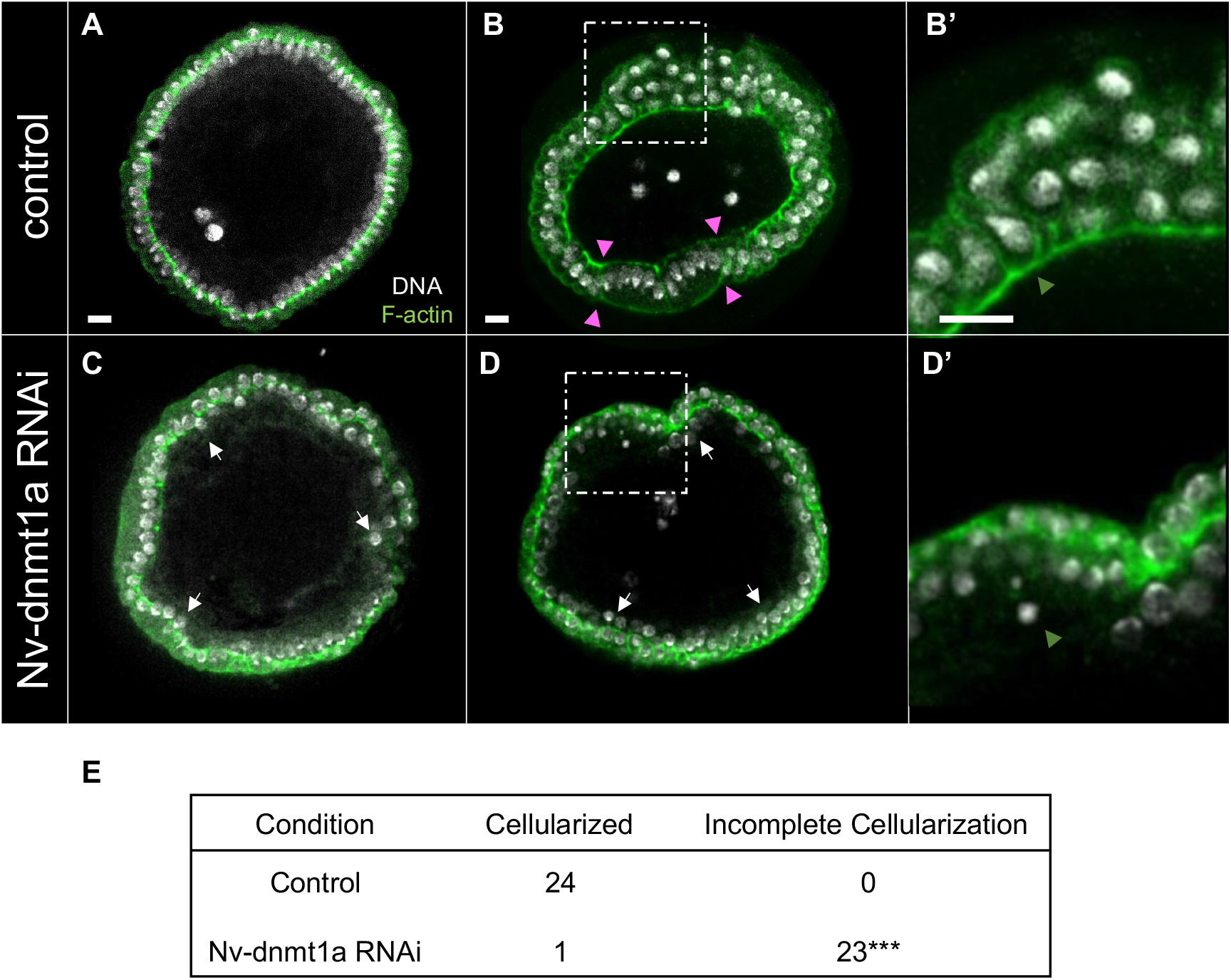
Transverse views of control and *Nv-Dnmt1a* RNAi embryos during cellularization and at the onset of gastrulation. Green, phalloidin (F-actin). white, DAPI (DNA). (A) Control embryo during mid cellularization. (B) Control embryo at the onset of gastrulation. Pink arrowheads demarcate the borders of the presumptive mesoderm. (B’) Zoomed in view of fully cellularized control embryo. Orange arrowhead points to the basal actin cable. (C) *Nv-Dnmt1a* RNAi embryo during mid-cellularization. White arrows point to examples of disordered nuclei. (D) *Nv-Dnmt1a* RNAi embryo at the onset of gastrulation. White arrows point to examples of non-cellularized, disordered nuclei. (D’) Zoomed in view of an *Nv-Dnmt1a* RNAi embryo that fails to fully cellularize. Orange arrowhead points to an non-cellularized nucleus (note the absence of the basal actin cable). (E) Table with n values of cellularized and non-cellularized control and *Nv-Dnmt1a* RNAi embryos. *** = p value < 0.0001 determined by Fisher’s Exact Test. All scale bars (white lines in A-A’’’) represent 10 μm.

All (24/24) control embryos initiating morphogenesis had nuclei that were fully enveloped by the plasm membrane and had an actin cable along the basal (interior) surface of the cellular blastoderm (Fig. 2B-B’, orange arrowhead in 2B’ marks the basal actin cable). Enlarged, round serosal cells were present on the dorsal side, and the distinct population of presumptive mesodermal cells was present on the ventral side (Fig. 2C, pink arrowheads mark the borders of the presumptive mesoderm). In contrast, *Nv-Dnmt1a* pRNAi embryos had many unencapsulated nuclei, and lacked the cable of actin along the interior surface of the blastoderm (Fig. 2D-2D’, orange arrowhead marks non-cellularized nuclei and obvious absence of the basal actin cable), consistent with failed cellularization (23 out of 24 pRNAi embryos were incompletely cellularized). In addition, the blastoderm was multilayered at many locations, and clearly distinct serosal and mesodermal precursors were not observed (Fig. 2D).

### Nv-Dnmt1a is required for global gene body methylation

The failure of cellularization and morphogenesis are widely conserved indicators of disrupted MZT (38–40). We hypothesized that *Nv-Dnmt1a* is required for proper regulation of the MZT, and its reduction by pRNAi would abrogate DNA methylation and consequently disrupt zygotic genome activation in the early embryo of *N. vitripennis*. We thus sought to characterize the DNA methylation patterns in the early embryo of *N. vitripennis* and to examine how this pattern was affected when *Nv-Dnmt1a* was depleted by pRNAi.

In control embryos collected at the late blastoderm stage (7-9 hours after egg-lay) we identified 151,480 methylated CpGs (mCGs) in a union set representing all of the different mCGs across three biological replicates. Each sample contained similar numbers of mCGs across the three biological replicates (ranging from 136,410 to 136,717). The vast majority of these mCGs were found within gene bodies (Table 1).

**Table 1.** Summary CpG statistics and non-conversion rates for each sample.

| Sample | Non-conversion rate | Median CpG coverage | Mean CpG coverage | Methylated CpGs (mCGs) | mCGs within gene bodies |
| --- | --- | --- | --- | --- | --- |
| Control 1 | 0.0052 | 14 | 14.92 | 136,600 | 126,333 |
| Control 2 | 0.0051 | 15 | 15.63 | 136,410 | 126,174 |
| Control 3 | 0.0055 | 14 | 14.75 | 136,717 | 126,364 |
| RNAi 1 | 0.0052 | 14 | 14.30 | 98,511 | 91,874 |
| RNAi 2 | 0.0051 | 14 | 14.48 | 8,017 | 7,720 |
| RNAi 3 | 0.0053 | 14 | 14.91 | 15,305 | 14,669 |

Mean fractional DNA methylation of gene bodies followed the canonical bimodal distribution (Fig. 3A) as seen in other invertebrate species including insects (21). Following the criteria used in a previous study, we designated genes with gene body fractional methylation levels of at least 0.02 as methylated genes (21). By this criterion we found 5,423 methylated genes in the blastoderm embryo, where 99% of our methylated genes in the embryo overlapped with methylated genes identified in adult *N. vitripennis* tissues (30), where a slightly different method to define methylated genes was used. This indicates that our results are highly compatible with those of Wang et al. (30), and are consistent with the idea that GBM patterns are generally stable across life stages.

**Figure 3.**
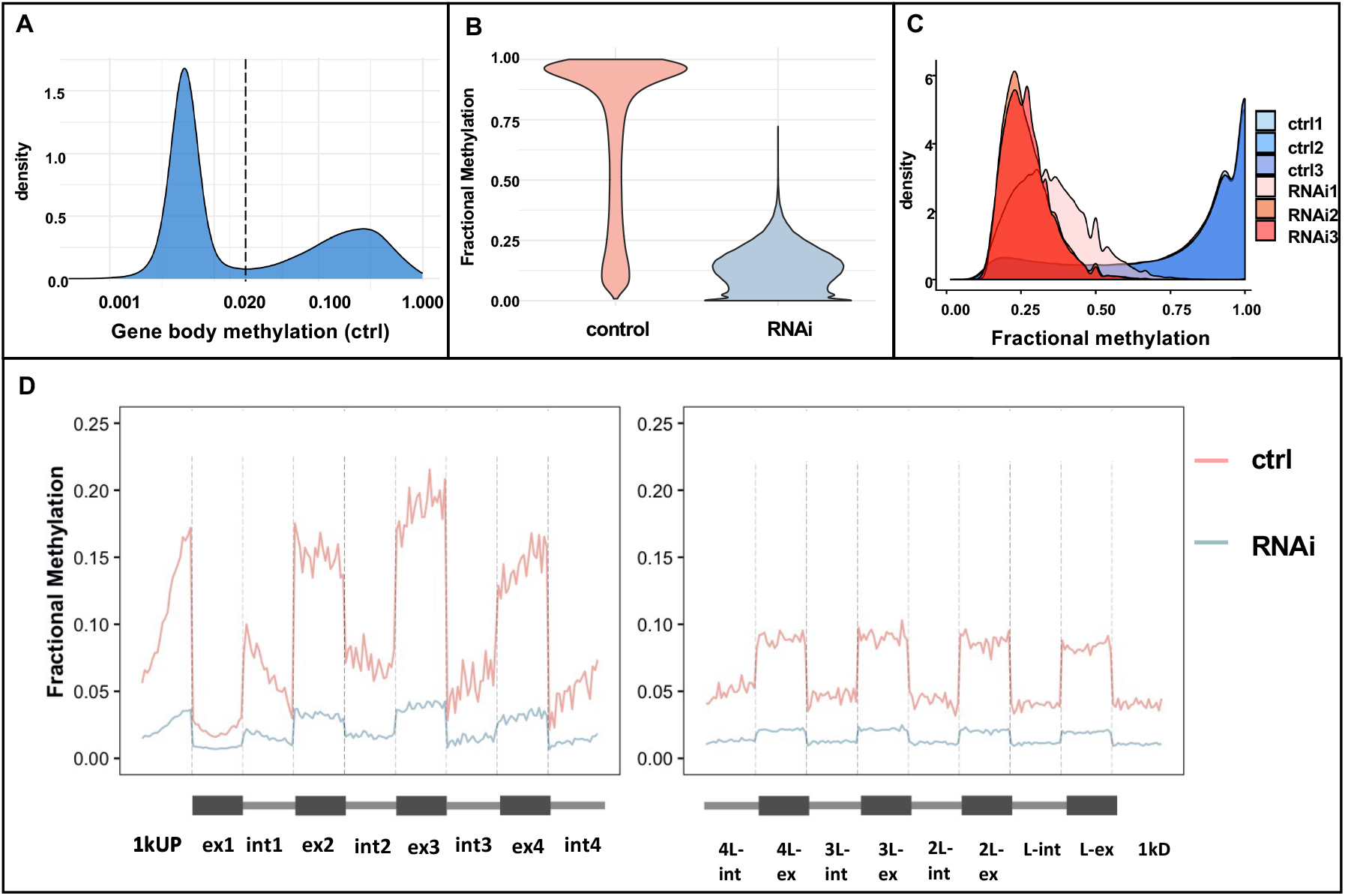
Global DNA methylation in control and *Nv-Dnmt1a* pRNAi late blastoderm *N. vitripennis* embryos. (A) An example of distribution of fractional DNA methylation in control embryo exhibiting the classical bimodal distribution. We defined methylated genes as genes with a minimum of 0.02 gene body methylation (shown as the dashed line) to classify methylated and unmethylated genes. (B) DNA methylation is substantially reduced in *Nv-Dnmt1a* pRNAi samples compared to the control samples. (C) Distributions of fractional methylation of CpGs that are methylated at least one control sample demonstrate that CpGs lose DNA methylation in RNAi embryos. (D) Distribution of mean fractional methylation in the first and last four exons and introns of methylated genes. Control samples shown in red, RNAi embryos shown in blue.

We knocked-down *Nv-Dnmt1a* mRNA with pRNAi and examined DNA methylation at the same developmental stage (7-9 hours after egg lay) as the control. Based on our RNA-seq results, *Nv-Dnmt1a* mRNA levels were reduced by 5-fold or more in late blastoderm pRNAi samples relative to late blastoderm control samples (41). We observed a significant reduction in DNA methylation at 99.7% of methylated CpGs in the genome in pRNAi embryos. The magnitude of the reduction was substantial, as we found an average of 81.5% DNA methylation depletion per mCG (Fig. 3B, 3C and Table 1).

While DNA methylation was strongly reduced genome-wide, the intragenic pattern of GBM was maintained. For example, mCGs were enriched in exons and were more numerous at the 5’ end of the coding region of gene bodies in control embryos (Fig. 3D) similar to patterns found in other insects and other developmental stages (29, 42, 43). This pattern was maintained in pRNAi embryos, despite the drastic overall reduction in DNA methylation levels (Fig. 3D). This suggests that *Nv-Dnmt1a* is required for gene body methylation but does not govern the 5′ and exon-biased pattern of DNA methylation that we see in *Nasonia* and other hymenopteran insects.

### Loss of Nv-Dnmt1a function disrupts the regulation of a large proportion of the early embryonic transcriptome

Given the profound effects of *Nv-Dnmt1a* on global DNA methylation and early developmental events, we sought to understand the effects of *Nv-Dnmt1a* knockdown on gene expression over time in the early embryo. Although the developmental effects *Nv-Dnmt1a* pRNAi were not visible until cellularization and the onset of morphogenesis, we hypothesized that changes in the transcriptome in response to disrupted development would occur earlier in development, leading up to the multifaceted failure of development at gastrulation.

To test this, we performed RNA-seq on timed egg collections from mock injected (control) and *Nv-Dnmt1a* dsRNA injected wasps at time points that cover early embryogenesis from egg laying to gastrulation. The collection time points were: freshly laid eggs (0-2 hours after egg lay), early blastoderm (3-5 hours after egg lay), middle blastoderm (5-7 hours after egg lay) and late blastoderm/onset of gastrulation (7-9 hours after egg lay).

We found 835 and 888 differentially expressed genes (DEGs) in the egg and early blastoderm stages, respectively. The affected loci at these stages likely reflect the effects of the knockdown in the maternal nurse cells, as there is little to no zygotic transcription at these stages. These effects were mild in both the number of genes affected and the magnitude of the effects compared to the following two stages (Fig. 4A). In the middle-blastoderm stage, where the zygotic genome is first broadly activated and maternal mRNAs are being cleared, 2744 genes showed significant differential expression, while in the late blastoderm stage 1831 were significantly differentially expressed.

**Figure 4.**
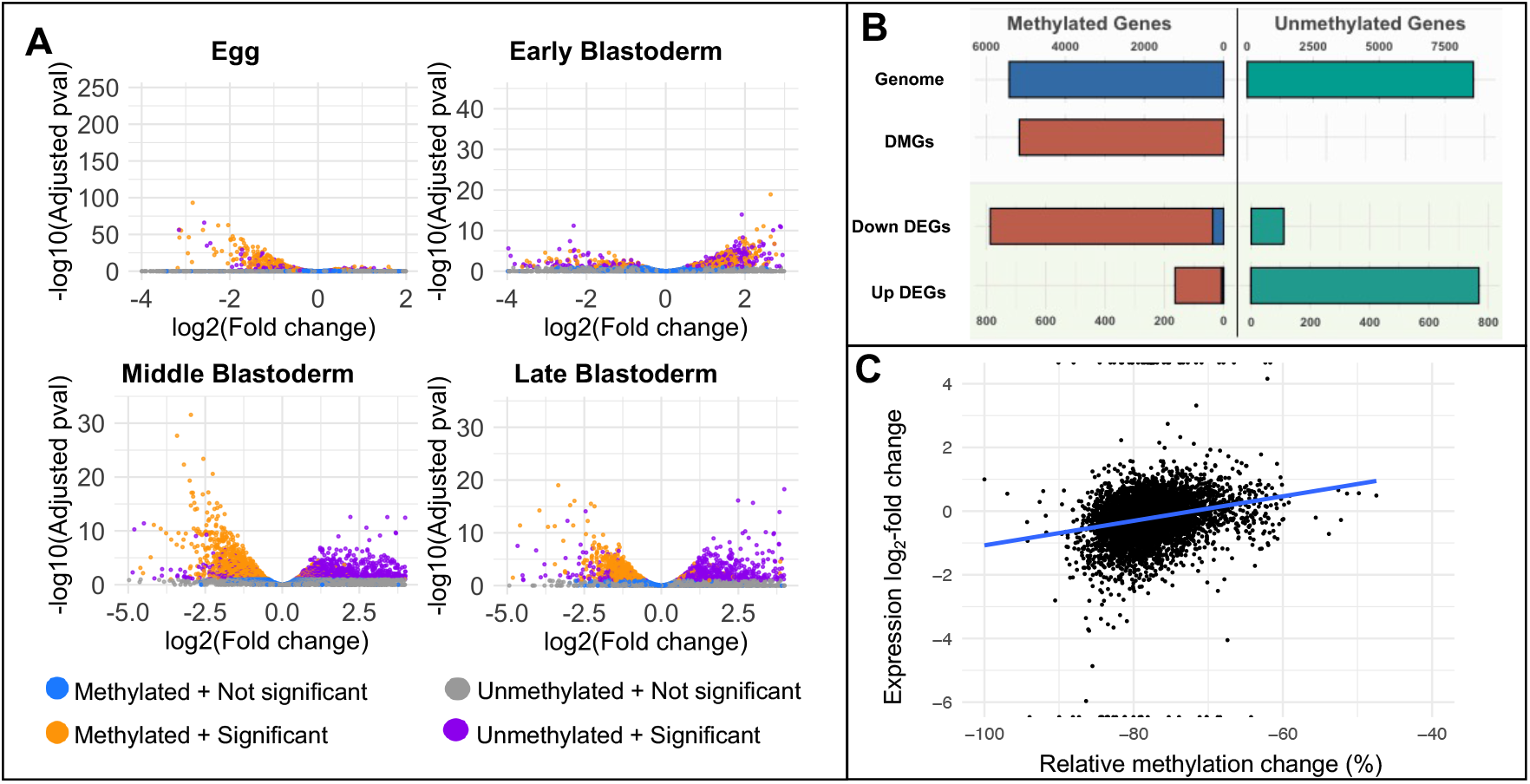
Gene expression of methylated and unmethylated genes in control and *Nv-Dnmt1a* RNAi *N. vitripennis* embryos. (A) Volcano plots showing differentially expressed genes between control and *Nv-Dnmt1a* RNAi in embryos of different developmental stages. X axis is log_2_ fold change in *Nv-Dnmt1a* RNAi samples relative to control samples. Y axis is log_10_ adjusted p-values. Golden dots are methylated, significantly differentially expressed (FDR-corrected P ≤ 0.1). Purple dots are significant and DEGs that are unmethylated (FDR-corrected P ≤ 0.1). Blue and grey are non-significant and represent methylated and unmethylated genes, respectively. (B) Distribution of methylated and unmethylated genes for all genes in the genome, differentially methylated genes (DMGs), down-regulated differentially expressed genes (Down DEGs), and up-regulated differentially expressed genes (Up DEGs). (C) The correlation between relative GBM change and relative expression change (calculated as log_2_-fold change) in the RNAi samples compared to the control samples was positive and statistically significant (Spearman’s rank correlation coefficient = 0.37, P < 2.2e-16).

Aside from the massive jump in the number differentially expressed genes at the mid-blastoderm stage, other intriguing patterns emerge at this time. Roughly equal numbers of the DEGs were up-regulated as were down-regulated. However, there was a clear bias in the methylation status of the DEGs and their direction of expression change. For example, 1201 out of 1331 (90%) of the down-regulated transcripts were derived from methylated genes in the middle blastoderm stage, while 1227/1413 (87%) upregulated transcripts came from unmethylated genes (Fig. 4A). Both of these observations were statistically significant (Fisher’s exact test, P < 0.001). A nearly identical pattern was maintained into the late blastoderm stage (Fig. 4A, B).

The strong over-representation of methylated genes among down-regulated genes following *Nv-Dnmt1a* pRNAi fits well with previous observations in insects connecting DNA methylation with high and stable gene expression (30, 32, 44). Furthermore, we have found that the magnitude of the reduction in expression was significantly correlated with the degree of lost methylation after pRNAi (Fig. 4C, Spearman’s rank correlation coefficient = 0.37, P < 2.2e-16). These observations imply that GBM plays an important role in maintaining normal levels of high expression from methylated loci, and its loss leads to widespread reduction in transcription from these loci. Assuming that GBM is the major direct output of *Nv-Dnmt1a* function, the effects on unmethylated DEGs (uDEGs), and potentially some methylated DEGs, may be due to indirect effects downstream of the initial reduction of mDEGs after pRNAi.

While indirect effects on unmethylated genes could be predicted, the very strong bias toward upregulation of this gene class was quite surprising. We compared our raw RNA-seq data and DESeq2 normalized data, to investigate if any technical bias due to the normalization method could have affected our inference of gene expression changes. There were no concerning or obvious differences between the distribution of raw read counts and the normalized counts used for our analyses (*SI Appendix*, SI Fig. 2). Further, the size-factors required for normalization were all relatively small (Supplementary Table 1).

### Methylated and unmethylated genes show stark differences in their regulation during zygotic genome activation

We hypothesized that unmethylated genes may be upregulated due to indirect effects of gene expression changes of methylated genes. Thus, we further examined all methylated and unmethylated genes over developmental time, focusing on those that showed differential expression between control and *Nv-Dnmt1a* pRNAi treatments in at least one stage (a total of 3904 genes).

We identified 1,769 methylated differentially expressed genes (mDEGs). Plotting the expression levels of the mDEGs revealed consistent unimodal distributions with peaks centered near the median expression level in each stage (Fig. 5). A similar pattern is seen when all methylated genes are plotted (*SI Appendix*, SI Fig. 4). *Nv-Dnmt1a* pRNAi did not affect the shape of these distributions, but rather resulted in a modest, but significant shift of the distributions downward (Fig. 5). This is consistent with *Nv-Dnmt1a* dependent GBM playing a role in aiding the efficiency of transcription of methylated genes.

**Figure 5.**
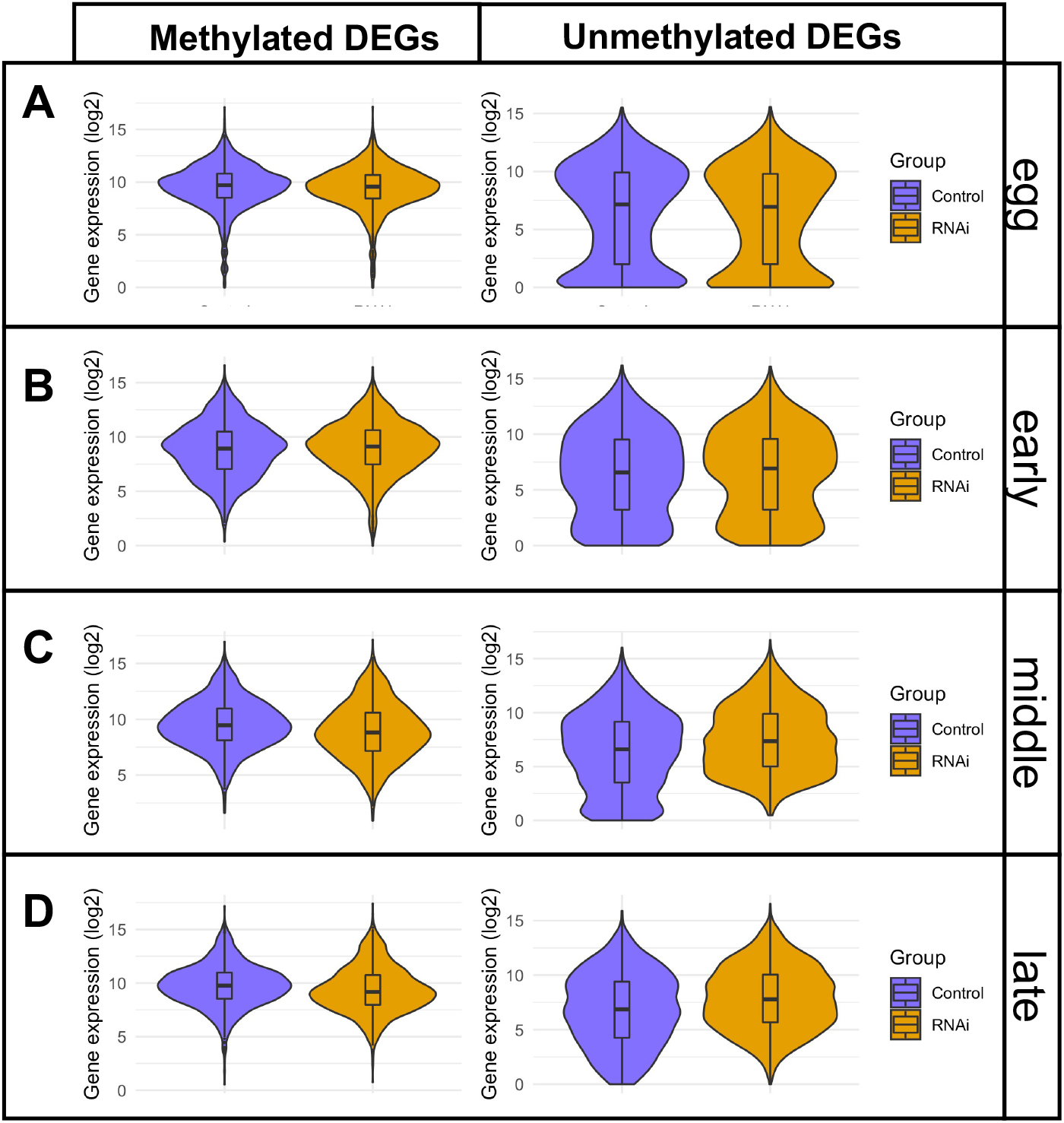
Violin plots of methylated and unmethylated differentially expressed genes (DEGs) in control and *Nv-Dnmt1a* pRNAi eggs, early blastoderm embryos, middle blastoderm embryos, and late blastoderm embryos.

We identified a total 2135 unmethylated DEGs (uDEGs). The distribution of expression levels for these genes was more complicated, showing a bimodal tendency, with a peak at very low expression levels during the egg through middle blastoderm stages (Fig. 5A-C), and less pronounced peaks at higher levels that did not overlap the medians. *Nv-Dnmt1a* pRNAi had little effect on global distribution of uDEG expression levels in the egg and early blastoderm stages (Fig. 5A, B), consistent with the minor effects of pRNAi on these stages reported earlier (Fig. 4A). However, in the middle and late blastoderm stages (Fig. 5C, D), the median uDEG expression increases slightly in pRNAi samples compared to control embryos, and there was a pronounced reduction of the lowest expression levels (below 2.5). This indicates that uDEGs that were effectively absent in control embryos were present at meaningful levels after *Nv-Dnmt1a* pRNAi. We propose that failure to either clear maternal RNA or repress transcription of these factors plays a crucial role in developmental failure at cellularization.

### Nv-Dnmt1a is required for the expression of housekeeping genes and mRNA decay factors

We examined the annotations of the DEGs and identified many differentially expressed factors, such as RNA binding and regulating factors (including the well I known MZT factors *Smaug* and the CCR-NOT deadenlyase complex), transcription factors, cyclins and cytoskeletal components that could plausibly play important roles in producing the observed patterns of gene expression change and developmental failure after *Nv-Dnmt1a* pRNAi. None of these candidates were either completely reduced, nor highly upregulated, so neither loss of function nor gain of function experiments would likely produce phenocopies of the *Nv-Dnmt1a* pRNAi. Rather, the observed patterns are consistent with moderate disruption of several genes interacting to produce the complex cellularization and morphogenesis phenotypes.

Gene ontology analyses found that terms involved in metabolism, and other basal cell functions were the most significantly and consistently enriched in both in methylated and unmethylated DEGs throughout development (*SI Appendix*, SI Table 2). Interestingly, RNA degradation is an enriched category of down-regulated methylated genes, (*SI Appendix*, SI Table 2), consistent with our RNA-seq results.

## Discussion

A previous study found that one of the DNMT1 orthologs in the wasp (*Nv-Dnmt1a*) was required for *Nasonia* embryogenesis, while the other two DNMT1 paralogs (*Nv-Dnmt1b, Nv-Dnmt1c*) and the DNMT3 ortholog *Nv-dnmt3* were dispensable for embryonic development (35). *Nv-Dnmt1a* pRNAi embryos failed to form segments and the knockdown was lethal (35) which led us to further examine the phenotype in more detail (45, 46).

We found that the first signs of developmental failure are nuclei that fail to be correctly cellularized. This directly precedes failure of the morphogenetic processes of mesoderm internalization, and migration of the serosa. These are all developmental events that are strongly affected when the MZT is disrupted in *N. vitripennis* and *D. melanogaster*. We hypothesized that *Nv-Dnmt1a* is required for GBM and the regulation of gene expression during the MZT. Consistent with this, *Nv-Dnmt1a* knockdown resulted in major disruption of gene expression levels during early embryogenesis. The most significant alterations to the transcriptome occurred at the mid blastoderm stage, which correlates well with when the major wave of zygotic genome activation occurs in *N. vitripennis*. This observation, in combination with our finding that GBM was significantly reduced after *Nv-Dnmt1a* pRNAi, strongly indicates that loss of gene body methylation disrupts the MZT in *N. vitripennis*.

The exact mechanism by which *Nv-Dnmt1a* regulates the MZT and downstream processes is not yet understood. However, a crucial observation was that there was a very strong correlation between gene body methylation state and the direction of expression change after *Nv-Dnmt1a* pRNAi: downregulated genes in pRNAi embryos were almost exclusively methylated genes (which were confirmed to have reduced GBM), and upregulated genes were conversely almost exclusively unmethylated (Fig. 4).

The correlation of reduced GBM with reduced gene expression among methylated genes after *Nv-Dnmt1a* pRNAi is consistent with previous observations that GBM is associated with high, steady levels of gene expression (30, 47). However, it has not been conclusively shown that GBM directly increases levels of gene expression, and the exact role played by GBM has been controversial (48, 49). Comparison of orthologous gene expression between plants species or populations that had either maintained or lost GBM capability revealed little or no effect of GBM on expression levels of ancestrally methylated genes, indicating that GBM may have no appreciable effect on transcription (48–50). In insects, knockdown of DNMT1 orthologs lead to the loss GBM in the roach *Blatella* and the milkweed bug *Oncopeltus* (29, 51). In the roach, loss of DNMT1 led to disruption of methylated gene expression levels, but no bias toward reduced expression was observed, while in the milkweed bug no change in gene expression levels was observed for any genes (29, 51).

The above results appear to conflict with our findings that, after *Nv-Dnmt1a* pRNAi, the vast majority of methylated genes that are differentially expressed show reduced expression, and that the magnitude of reduced expression directly correlated with the degree of gene body methylation loss. This may indicate that the role of GBM methylation is quite labile in evolution, and its significance and exact role might vary from species to species. We also propose that our highly focused analysis of early embryogenesis is advantageous for detecting moderate and transient effects of altered GBM. The pre-gastrulation embryo is a relatively homogenous tissue, where cells have not completed differentiation. Given that we are examining the very early stages of embryo development, we might be able to capture early response to the loss of GBM without being obscured by noise introduced by indirect effects. In the other studies, gene expression was examined in complex tissues that were developed well beyond the earliest potential effects of GBM loss. We believe further taxonomic sampling that focuses on time limited and homogenous tissues will reveal the relative importance of these ideas.

While our data strongly suggests that Dnmt1a regulates gene expression through GBM in *Nasonia*, there is evidence that DNMT1 orthologs have roles independent of their methylation functions in other insects. *Dnmt1* knockdown data in milkweed bug *Oncopeltus fasciatus* and the beetle *Tribolium castaneum* suggest that Dnmt1 orthologs may have molecular functions unrelated to their methylase activity (29, 52, 53). We cannot exclude that a non-methylating function of *Nv-Dnmt1a* exists, and that it could contribute to the complex phenotypes we observed after knocking down *Nv-Dnmt1a* with pRNAi. However, the strong correlation of GBM loss and reduced methylated gene expression after Nv-Dnmt1a knockdown is most easily explained by an important role for the canonical methylase activity of *Nv-Dnmt1a* in the embryo.

## Conclusion

The reduction of GBM and methylated gene expression after pRNAi was also associated with major developmental defects in embryogenesis, providing strong evidence that *Dnmt1a* and GBM are essential components for the MZT in *Nasonia*. We have shown that pRNAi results in incomplete cellularization and gastrulation failure, which is similar to what has been observed when we inhibit zygotic transcription or knockdown essential MZT genes, such as Smaug or Zelda. Importantly, we show that GBM loss is significantly associated with a reduction in methylated gene expression during zygotic genome activation. Our results further suggest that the indirect effects of *Nv-Dnmt1a* pRNAi are mediated by both disruption of mRNA degradation, and misregulation of transcription from the zygotic genome. These data provide the first evidence that GBM is essential for both the full activation of methylated genes and the subsequent proper regulation of unmethylated genes during the maternal-zygotic transition. The exact mechanisms by which GBM affects transcription, and how disruption of GBM leads to the failure of developmental regulatory networks in *Nasonia* will be important questions to address in order to understand the function and evolution of this widespread but poorly understood epigenetic modification.

## Materials and Methods

### Live imaging

All embryos were live-imaged as described in Arsala & Lynch 2017 (38). Briefly, we live-imaged all embryos at 28C for 15 hours on an Olympus BX-80 inverted microscope under 30X silicone immersion and DIC optics.

### Phalloidin staining & transverse sectioning

Control injected and *Nv-Dnmt1a* dsRNA (1ug/ul) injected mothers were allowed to lay eggs on hosts for 2 hours at 25C. Freshly laid eggs were collected from hosts and aged on 1% agarose PBS plates at 25C until they were 8-10 hours old, so that all embryos were aged to correspond with the onset of gastrulation. We hand dissected the chorions of fixed embryos. Afterward, the fixed embryos were placed in 1X PBS and stained with AlexaFluor 488 phalloidin at a 1:250 dilution. The stained, fixed embryos were placed in vectashield with DAPI overnight. Finally, the embryos were laid flat on a cover slip and were cut in half along the transverse axis using a razor slotted into an embryo ‘guillotine.’ The resulting embryo ‘halves’ were flipped ‘up’ on the cover slip and imaged on an Andor Revolution WD spinning disc confocal system. The samples were illuminated with 488nm (phalloidin) and 405nm (DAPI) diode laser and Z-stack images were captured using a 30x objective at 1micron increments.

### RNA extraction

Embryos were collected and aged as described above. We designed 3 non-overlapping dsRNA constructs (see *SI Appendix*, Table 3 for primer information) and injected 100 females per biological replicate with either 1μg/mL dsRNA diluted in water or with water as a control. The females were left unmated, so that all resulting progeny were male as the variation in sex and ploidy could confound our analysis. The injected females were allowed to lay eggs that were aged at 25C until they reached the following four time points across the MZT: 0-2hours to generate a freshly laid egg transcriptome, 3-5hours to generate an early blastoderm transcriptome, 5-7 hours to generate a middle blastoderm transcriptome, 7-9 hours to generate late blastoderm embryos that are at the onset of gastrulation. Once the embryos reached the time points, we isolated 100ng of total RNA using TRI Reagent for all samples in biological triplicate. We confirmed the RIN value for all samples were above 9.0 using a 2100 Agilent Bioanalyzer and immediately prepared 24 libraries for sequencing.

### RNA library preparation and bulk RNA-sequencing

Using the 100ng of total RNA from each sample isolated above, we performed poly-A tail selection using the NEBNext Poly (A) mRNA magnetic isolation module (E7490) and immediately proceeded with library preparation. Libraries were prepared for bulk RNA-sequencing using the NEBNext Ultra Directional RNA Library Prep Kit for Illumina (E7420). The libraries were purified, validated and pooled before sequencing. The libraries were subjected to 100bp paired-end sequencing across 2 lanes on a HiSeq4000. Raw data from this analysis can be found in the NCBI BioProject archive with accession number PRJNA701367.

### RNA-seq analysis

To assess sequencing quality, we used FastQC and saw high quality reads across all samples. We trimmed adapters and removed low quality reads using Trim Galore-0.6.0 only using the --paired parameter. We reassessed the trimmed read quality using FastQC and proceeded to map the reads to genome using HISAT2-2.0.5 (hisat2 -p 8 --max-intronlen 10000 -q –x). Differential expression analysis was performed using the R package DESeq2(54). We filtered out lowly expressed genes (maximum read value of 10) from our analysis.

### DNA extraction and bisulfite library preparation

We extracted and purified genomic DNA from embryos using the QIAamp DNA Micro Kit (Qiagen 56304). 40ng of genomic DNA was bisulfite converted and libraries were prepared at the University of Chicago Genomics Facility using the Accel-NGS Methyl-Seq DNA Library Kit from Swift Biosciences. Raw bisulfite sequencing reads can be found in the NCBI BioProject archive with accession number PRJNA701143.

### Analysis of Bisulfite-sequencing data

CpG sites were determined to methylated using a binomial test (30, 31) where the rate of deamination representing the probability of success and total number of reads representing the number of trials (30, 31). P-values were then corrected for multiple testing (55) and CpGs with adjusted P-values of < 0.05 were labeled as methylated CpGs. Genes containing at least one methylated CpG were subsequently labeled as methylated genes. Fractional methylation of CpGs was calculated by dividing the number of methylated reads by the total number of reads and the fractional methylation of genes was calculated by taking the average fractional methylation of all CpGs within the gene body. For the differential methylation analysis, only CpG sites that were methylated in at least one control sample were retained similar to previous insect studies (44, 56). The DSS package (57) was used to assess differentially methylated CpGs with RNAi status as the sole predictor of methylation. P-values for each CpG site was adjusted for multiple testing (55) with significance level set at an adjusted P-value of < 0.1. Genes with at least one differentially methylated CpG were considered as differentially methylated genes.

## Author Contributions

D.A.: Conceptualization; Formal analysis; Supervision; Funding acquisition; Investigation; Visualization; Methodology; Writing - original draft; Writing - review and editing. X.W.: Conceptualization; Formal analysis; Investigation; Visualization; Methodology; Writing - original draft; Writing - review and editing. S.V.Y.: Formal analysis; Supervision; Visualization; Methodology; Writing - original draft; Writing - review and editing. J.A.L.: Conceptualization; Formal analysis; Supervision; Funding acquisition; Writing - original draft; Writing - review and editing

## Acknowledgments

We thank The University of Chicago Genomics Facility (RRID:SCR_019196), for their very valuable assistance with generating the WGBS and RNA-seq data. This work was supported by grant MCB1615664 from NSF to SVY, and by grants R01GM129153 and R03HD087476 from NIH to JAL.

## Supplementary Information for

**Fig. S1.**
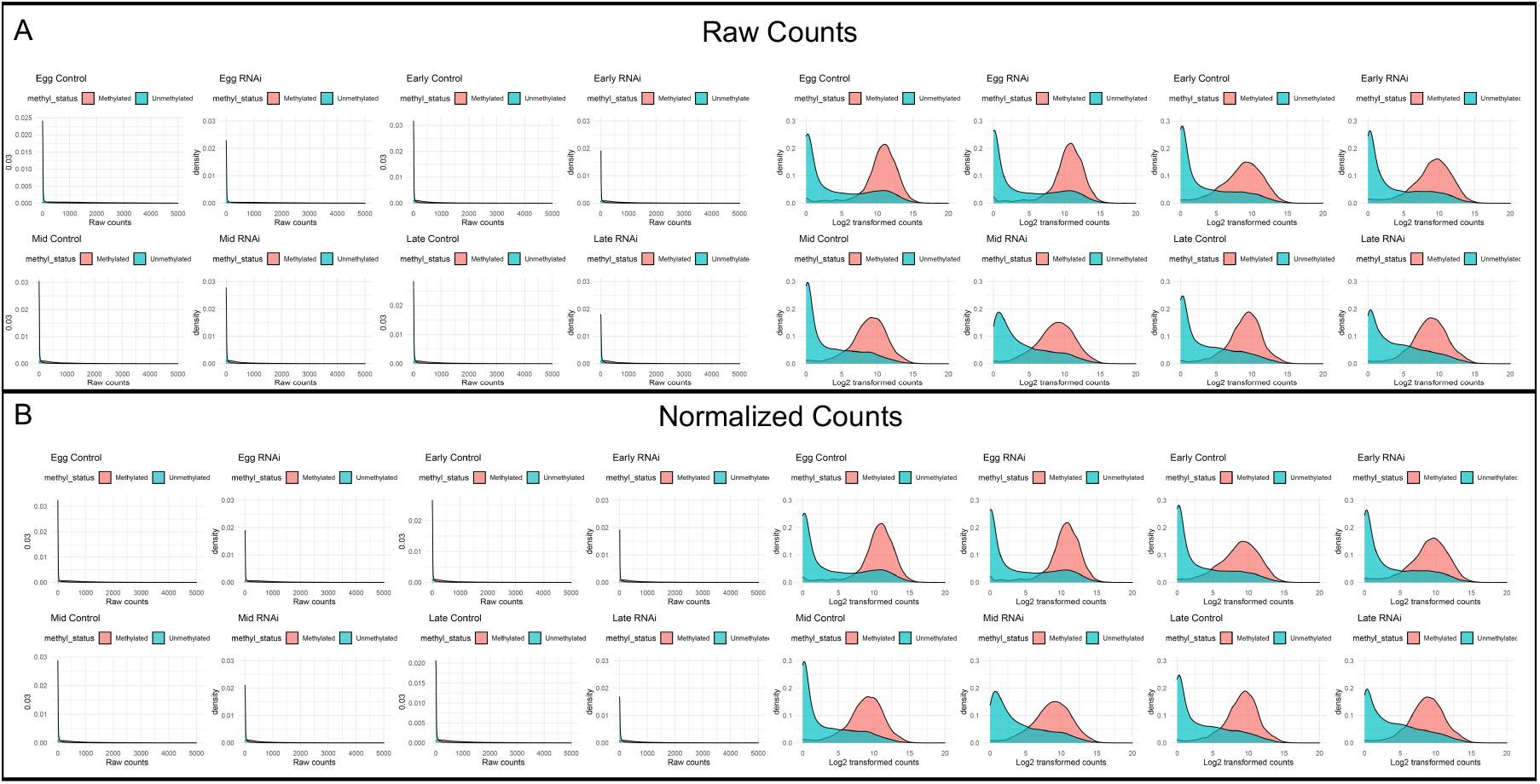
Density distributions of (A) raw transcriptome read 745 counts and (B) normalized counts from DESeq2. Data are log2-746 transformed. Methylated and unmethylated genes exhibit distinctive patterns of 747 expression consistent with previous studies. The density distributions of the raw 748 and normalized counts are highly similar.

**Fig. S2.**
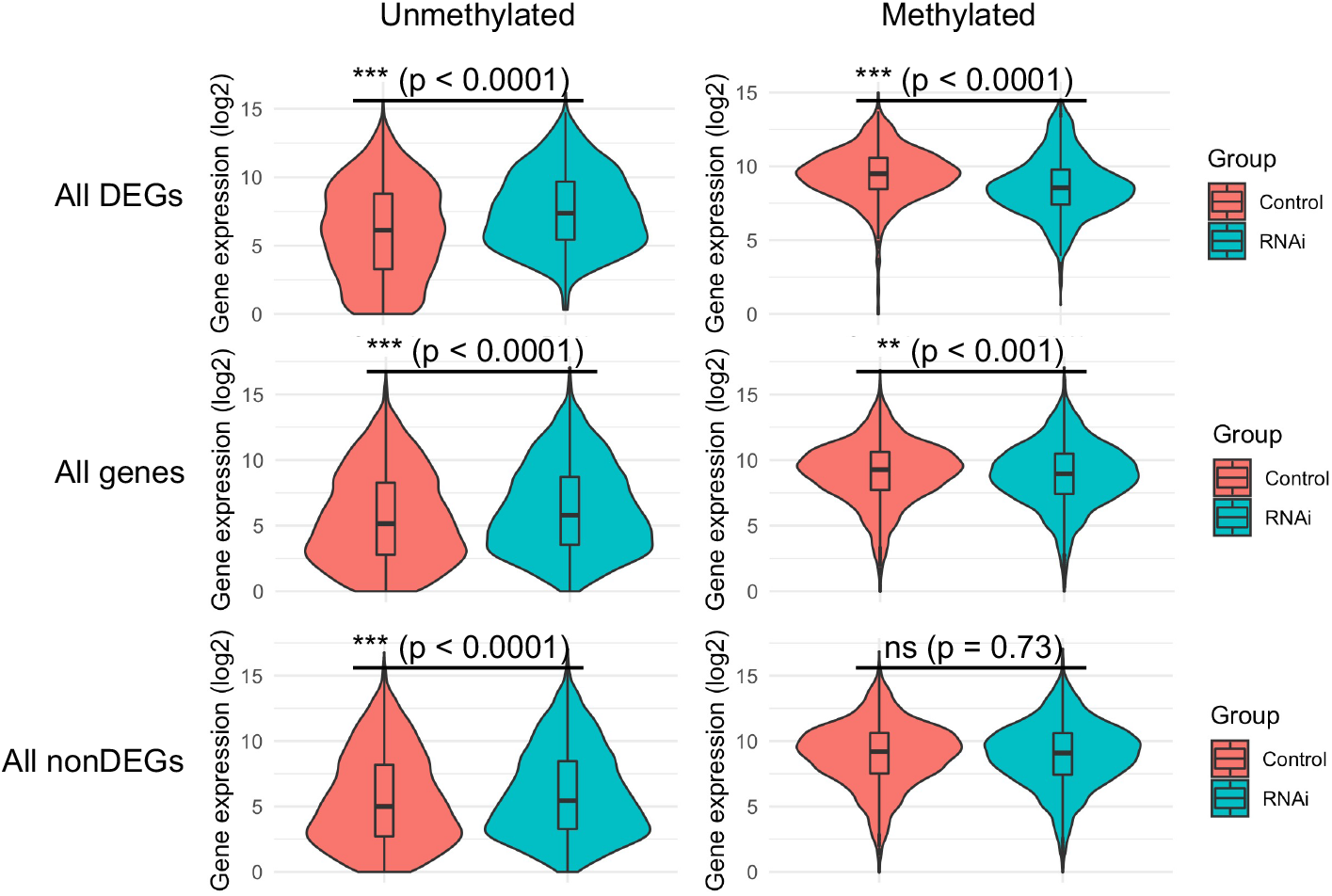
Violin plots of methylated and unmethylated genes during the late blastoderm stage in control and Nv-dnmt1a pRNAi embryos. Top row, all differentially expressed genes (All DEGs), middle, all genes, and bottom, all non-differentially expressed genes (All nonDEGs).

**Table S1.** Size Factors used for normalization of RNA-seq libraries.

| Sample | Size Factor |
| --- | --- |
| Egg control rep 1 | 2.008 |
| Egg control rep 2 | 2.618 |
| Egg control rep 3 | 2.389 |
| Egg RNAi rep 2 | 1.855 |
| Egg RNAi rep 3 | 2.422 |
| Early control rep 1 | 0.799 |
| Early control rep 2 | 0.789 |
| Early control rep 3 | 1.041 |
| Early RNAi rep 1 | 0.945 |
| Early RNAi rep 2 | 1.234 |
| Early RNAi rep 3 | 1.004 |
| Middle control rep 2 | 0.848 |
| Middle control rep 3 | 0.609 |
| Middle RNAi rep 1 | 0.709 |
| Middle RNAi rep 2 | 0.835 |
| Middle RNAi rep 3 | 0.843 |
| Late control rep 1 | 0.763 |
| Late control rep 2 | 0.891 |
| Late control rep 3 | 0.689 |
| Late RNAi rep 1 | 0.822 |
| Late RNAi rep 2 | 0.461 |
| Late RNAi rep 3 | 1.026 |

**Table S2.**
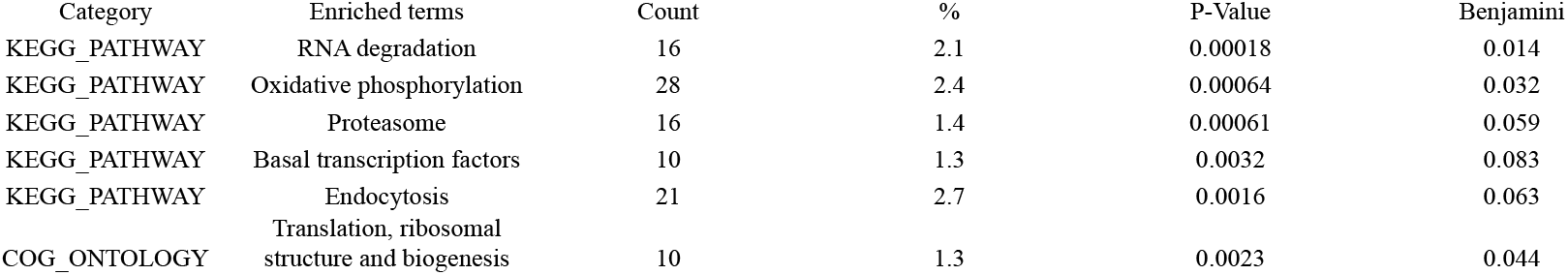
Gene Ontology analysis for down-regulated methylated genes in middle and late blastoderm Nv-Dnmt1a RNAi embryos.

**Table S3.**
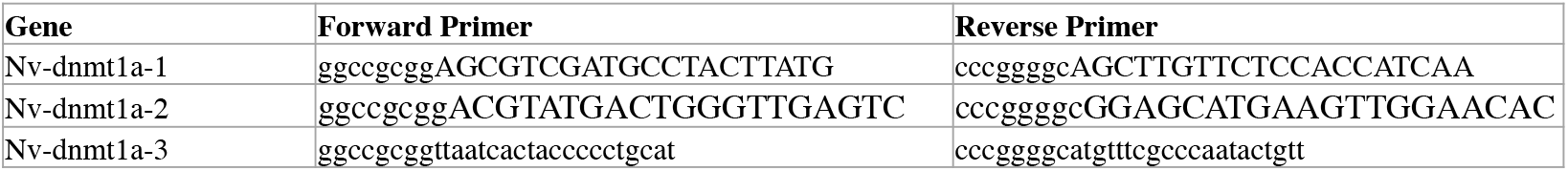
Primer sequences for *Nv-dnmt1a* knockdown.

